# Foot Sole Cutaneous Somatosensory Modulation Based on Balance Demands

**DOI:** 10.1101/2025.07.28.667302

**Authors:** Ali Doroodchi

## Abstract

Postural complexity may shape how the nervous system processes plantar cutaneous input. We tested whether somatosensory evoked potentials (SEPs) elicited by foot-sole stimulation scale with balance demands, hypothesizing larger responses for more complex tasks. Thirty-one healthy adults performed standing, straight-step, and diagonal-step conditions while receiving brief electrical stimulation to the stance foot sole; SEPs (P50, N90, peak-to-peak) were analyzed at Cz using pooled and order-specific approaches.

In the pooled analysis, peak-to-peak SEP amplitude was greater for both stepping conditions than standing (Standing vs. Straight, *P* = 0.041; Standing vs. Diagonal, *P* = 0.026). Order-specific analysis showed an early amplification: the first SEP (0.5 s after the warning cue) was larger for diagonal than straight stepping (peak-to-peak, *P* = 0.009; N90, *P* = 0.027). Source localization at N90 revealed greater activation during stepping than standing in paracentral gyrus & sulcus, inferior parietal angular gyrus, and superior parietal gyrus, consistent with enhanced sensorimotor processing under higher postural demands. Moreover, right paracentral gyrus & sulcus activity was higher for diagonal vs. straight stepping for the first and fourth SEPs.

Together, these findings indicate that increasing balance demands up-weight plantar afferent processing and recruit contralateral sensorimotor/parietal regions, particularly early in preparation, supporting the view that cortical sensory gain is tuned to postural complexity.

## Introduction

Safe, efficient movement depends on balance, which in turn requires postural control—the continuous regulation of body segments and postural muscle activity to keep the center of mass within the base of support. This chapter sets the foundation by outlining the determinants and sources of complexity in postural control from quiet standing to step initiation, then describing the cortical contributions that enable it: selective processing of task relevant sensory inputs (with emphasis on plantar cutaneous cues) and the feedforward planning of postural muscle activations that implement anticipatory postural adjustments. Together, these concepts frame the material that follows.

### Balance & Postural Control

To maintain balance, center of mass (CoM) must be maintained within the base of support (BoS) (Aruin et al., 2003; Lugade et al., 2011). Postural muscles aid in balance maintenance by modulating posture (Chang et al., 2010). For appropriate balance and postural control, postural muscles’ activation must be controlled, and appropriate sensory information must be utilized to base the postural muscular control upon (Danis et al., 1998).

### Postural Control Complexity

The complexity of postural control can be influenced by several factors, including the amount of torque each muscle must generate, the coordination required across muscle activation patterns, the precision of timing in these activations, and the level of balance threat that must be managed (Forbes et al., 2018; Horak, 2006). Requiring higher joint torques makes postural control more difficult because it amplifies neuromuscular noise, increases the need for agonist–antagonist coordination, quickens segment movements, and leaves the system with a smaller margin for error (Do Nascimento et al., 2005; Petrucci et al., 2018). More complex muscle activation patterns raise postural control demands because the nervous system must synchronize a larger, more overlapping set of muscular synergies within narrow time windows, a burden that grows with task difficulty and directly challenges balance stability (Ariani et al., 2022; Goodworth & Peterka, 2012). Tightening the time window for postural muscle activation makes balance more difficult because it eliminates feedback-correction time, so even slight timing errors can push the body outside its stability margin (Kawato, 1999). Greater displacement of the CoM relative to the BoS increases balance complexity because it narrows the margin of stability, leaving minimal room or time for corrective actions before a fall ensues (Leuthold & Schröter, 2011; Lugade et al., 2011).

### Anticipatory Postural Adjustments (APAs)

Initiating a step perturbs standing balance in two ways: unloading the swing foot instantly narrows the base of support to the stance foot alone, and the ensuing body thrust moves the CoM forward (Chen & King, 2024; Day & Bancroft, 2018). Anticipatory Postural Adjustments (APAs) are a set of postural modulations that occur prior to voluntary movements, aiming to maintain balance stability throughout the movement (Delafontaine et al., 2019; Farinelli et al., 2021). Shifts in the CoM during the APAs propels the body forward in the desired location, and it ensures balance is regained as landing the stepping foot re-establishes a stable BoS (Farinelli et al., 2021). These shifts in the CoM are influenced by the step’s characteristics such as its speed and direction (Bancroft & Day, 2016; Farinelli et al., 2021; Lyon & Day, 1997). Incorporating a diagonal impulse into the APA is more demanding than a straight-step APA because muscles must generate precisely scaled torques in both the sagittal and frontal planes, the multidirectional coordination tightens timing and synergy requirements, and the larger medio-lateral CoM shift relative to the stance foot BoS narrows the stability margin (Bancroft & Day, 2016; Delafontaine et al., 2019).

### Cortical Contributors

For appropriate postural control, appropriate sensory processing and postural muscle’s motor planning are vital (Forbes et al., 2018). Relevant sensory information must be processed effectively and integrated into the movement plan efficiently (Alfuth & Rosenbaum, 2012; Ariani et al., 2022). The movement plan must prepare the body for the desired task while also ensuring stability while performing it (Brown & Staines, 2015; Brunia et al., 2011). This section describes cortical mechanisms that may be involved in sensory processing and motor preparation required for balance and postural control.

#### Sensory Processing

Sensory information from a variety of modalities is sensed from various sensory organs. Irrelevant sensory information must be filtered, while relevant sensory information must be amplified and its processing enhanced to provide adequate information for basing the upcoming tasks upon (Mease et al., 2014; Sieveritz & Raghavan, 2021). Knowledge about an upcoming task presented prior to its execution directs attention towards sensory information relevant to the forthcoming task (ElShafei et al., 2018). Selective attention mechanisms enhance relevant sensory information such that motor preparation for an upcoming motor task can be done effectively. Anticipatory selective attention mechanisms allow this sensory modulation by influencing sensory filtering mechanisms occurring at the thalamus and affecting activation in appropriate sensory processing areas (Liu et al., 2016; Zhao et al., 2022). Enhanced relevant sensory information provides a high-fidelity platform for basing the motor task upon, improving motor preparation’s accuracy and efficiency (Brunia, 1993; Gaillard, 1977). With more relevant sensory information available, important factors in motor preparation, such as torque production, muscle activation timing and coordination can occur precisely (Goodworth & Peterka, 2012).

#### Sensory processing for Postural Control

Somatosensory information coming from the foot sole cutaneous information is vital for balance and postural control (Goodworth & Peterka, 2012; Kavounoudias et al., 1998). Foot sole cutaneous receptors directly measure body weight’s distribution across the BoS, which is vital in estimating the CoM and therefore maintaining balance (Kennedy & Inglis, 2002; Strzalkowski et al., 2018). Postural sways occur in directions opposite to foot sole tactile stimulations that activate foot sole cutaneous receptors but remain at an intensity below perceptual threshold (Kavounoudias et al., 1998). These postural sways mimic adjustments required for bringing CoM towards BoS center when it is naturally displaced outside BoS, but in this case, there was no actual CoM displacement to respond to (Kavounoudias et al., 1998). Furthermore, foot sole cutaneous information is amplified when preparing to take a step compared to standing (Fabre et al., 2023; Mouchnino et al., 2015). In sum, plantar cutaneous feedback is indispensable for balance maintenance because its continuous CoP encoding informs the CNS of CoM position relative to the BoS, enabling timely and appropriately scaled postural corrections.

### Electroencephalography (EEG)

Electroencephalography (EEG) measures the electrical dipoles generated from pyramidal neurons collective activation (Gable et al., 2022). If a sufficient number of pyramidal neurons in close proximity are activated simultaneously, their collective dipole can propagate through brain’s tissues via volume conduction and be measured via EEG electrodes placed on the scalp (Gable et al., 2022; Hádinger et al., 2022). Somatosensory evoked potentials (SEPs) are time-locked EEG responses to peripheral somatosensory stimulation, commonly used to assess cortical processing of somatosensory input (Kappenman & Luck, 2011). Their amplitude may provide information about the filtering applied to sensory information and processing intensity (Schubert et al., 2008).

Cortical activity’s source cannot be estimated from the surface EEG electrodes because the dipoles created from different neurons interact with each other, and the individual dipoles propagation are tempered as they go through different tissues such as the brain, meninges, and the skull (Gable et al., 2022). Source localization algorithms like exact Low Resolution Electromagnetic Tomography (eLORETA) aim to estimate the most probable location of cortical activity that may give rise to the recorded activity at the electrode level, by considering the attenuation imposed by different head tissues and the possible ways the dipoles may interact with each other (Faes et al., 2021; Jatoi et al., 2014). Therefore, eLORETA analysis may be used to explore SEP’s cortical origins more exhaustively than what may be interpretable from the data from EEG electrodes located on the scalp (Faes et al., 2021; Jatoi et al., 2014).

### Research Direction

Although the kinematics for different stepping movements and their proceeding APAs have been explored before, it remains unclear how postural complexity may influence task-relevant somatosensory processing. We aimed to investigate whether SEP amplitude differs with postural complexity, hypothesizing that greater postural complexity (e.g., in diagonal stepping) would amplify SEPs.

## Methods

### Sample

A total of 40 participants were recruited from the local population who met the following inclusion and exclusion criteria. Participants were included if they were between the ages of 18-40, and able to fluently communicate in English. Participants were excluded from this project if they had any current lower extremity musculoskeletal injuries/conditions (previous musculoskeletal injuries were accepted as long as recovered), lower extremity sensory injuries/conditions, neurological conditions that might affect gait or attention (such as Attention deficit hyperactivity disorder, Parkinson’s disease, stroke, etc.) circulatory problems in the lower extremity (e.g., Deep-vein thrombosis, impaired circulation), electronically active implants (e.g., pacemakers, cardiovascular defibrillators) or metallic implants in the foot.

Participants were recruited through the OurBrainsCan Registry and through a mass recruitment email from Western University’s registry. The University of Western Ontario’s research board committee approved this study. After enrolment, we were unable to book 5 participants’ data collection for this project. Of the remaining 35, 3 participants’ data were eliminated due to equipment technical difficulties, and 1 was eliminated due to deviations from the study protocol. 31 participants data are included in this project. The included sample size’s average age was 23.71 years (SD = 4.95), with 17 males and 14 females.

### EEG

The Biosemi Active 2 64-channel EEG system (Netherlands) was used with a sampling rate set to 1024 Hz. The 10-20 international montage system was used, and the electrode impedance was reduced to less than 10 microohms. Data collection was performed in an electromagnetic and acoustic shielded room.

### Tasks

EEG recording was done while participants performed three tasks: Standing, straight stepping, and diagonal stepping. For the standing condition block, data was recorded while participants stood still and received electrical stimulation (electrical stimulations paradigm is discussed shortly).

The Stepping tasks (straight and diagonal) were performed with electrical stimulation following auditory cues (auditory cues and electrical stimulation are discussed shortly).

These stepping tasks were selected based on established differences in their APA kinematics (Bancroft & Day, 2016; Lyon & Day, 1997). Diagonal stepping requires a more pronounced asymmetrical CoP shift to meet the greater balance perturbations of the diagonal step vector, demanding greater postural control compared to straight stepping for safe and efficient execution (Bancroft & Day, 2016; Lyon & Day, 1997). Quite standing imposes minimal threats to balance, requiring significantly smaller CoP and CoM control compared to the stepping tasks (Forbes et al., 2018; Hijmans et al., 2008). Consequently, differences in motor planning processes across these tasks may reflect increased sensorimotor integration and preparatory neural activity as postural demands intensify.

In the stepping conditions, participants were asked to step forward to a length that felt natural for initiating walking. For diagonal stepping trials, the step was performed approximately 45 degrees lateral from the forward direction. All Stepping trials were done with the right foot as the stepping foot, and the left foot as the stance foot. Participants were instructed to maintain stepping characteristics such as step length, speed, and angle, across trials as consistent as possible across trials. To minimize eye and head movement artifacts, a fixation point was placed at eye level on the wall directly in front of the participants, who were instructed to maintain their gaze on this point and keep head movements to a minimum.

#### Stepping Trial Quantities

40 steps per stepping type were performed, totaling 80 steps. The participants performed these steps in 2 blocks of 40 with a 5-minute rest in between, where participants were given the opportunity to sit down. The stepping type was randomized for each participant, until the maximum number of a step type was reached. Once the maximum number of a step type was reached (40 per block), the remainder of the step types were switched to the unfinished movement.

### Auditory Cues

Auditory cues, played through speakers, instructed the participants about the stepping type they should perform. The auditory cues included a ‘*Warning’* cue and a ‘*Go’* cue. The warning cue informed participants about the type of the upcoming stepping task (either straight or diagonal). The warning auditory cues were “One”, associated with forward stepping, and “Two”, associate with diagonal stepping. Two seconds after the termination of the Warning cue, a 440 Hz beep sound was played as the Go signal, instructing the participants to execute the stepping task. Participants were instructed to be prepared for the step after hearing the warning cue (i.e., “One” or “Two”) but not to initiate the step until they hear the imperative cue (the 440 Hz beep). The participants were instructed to maintain their position after stepping until they hear the return cue. An auditory cue “Return” was played 3s after the termination of the Go cue to instruct the participants to return to their original stance.

### Electrical stimulation

The Digitimer DS7HA (United Kingdom) was used for electrical stimulation. The anode was placed on the calcaneus and the cathode proximal to the metatarsal heads (both anode and cathode were on the foot sole), in line with the paradigms used in previous literature (Fabre et al., 2022, 2023; Lhomond et al., 2019; Mouchnino et al., 2015). Figure 1. shows electrodes’ location. The stimulation intensity was set to 25% above the participants’ perceptual threshold and was delivered at a 2 Hz frequency with a pulse width of 0.02 seconds, in line with the literature (Fabre et al., 2022, 2023; Lhomond et al., 2019; Mouchnino et al., 2015). To determine the perceptual threshold, the voltage was set to 200V and current set to 0 mA (minimum values possible on the device). The current was gradually increased until participants reported feeling the stimulation distinctly and consistently. Voltage remained unchanged at 200V. Then, the current was reduced to a level where participants could no longer feel the stimulation and then increased to a perceptible level. This process was carefully repeated multiple times until the lowest current where the participants could consistently feel the stimulation was found. This current was then multiplied by 1.25 to determine the current used in the study. All participants reported the stimulation to be comfortable and not painful. Figure 2. summarizes the electrical stimulation procedure and the timing of the cues.

**Figure 1:**
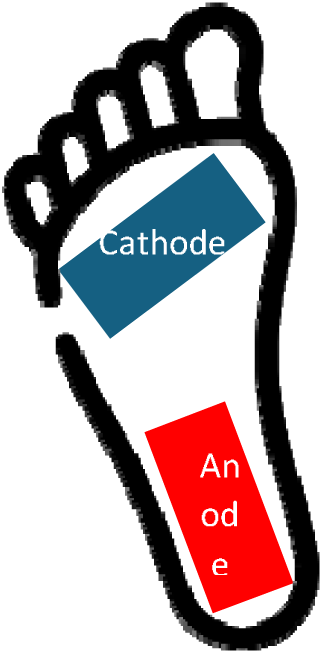
Stimulation Electrode Location on the Stance foot (left)

The electrical stimulations were ongoing throughout the blocks with electrical stimulations, but the simulations were set to occur at fixed times relative to the auditory cues. In Stepping trials, 4 electrical stimulations always occurred at 0.5s, 1s, 1.5s and 2s after the participants heard the warning stimuli but have not yet started the Stepping movement’s executions. These 4 electrical stimulations presented in a stationary preparatory period were used for SEP’s analysis for the stepping trials. With 40 steps per direction in the blocks with electrical stimulation, 160 electrical stimulations were delivered per stepping type. For the Standing block, 300 electrical stimulations following the same stimulation procedure were delivered. The auditory cues were not presented in the Standing block to prevent contamination from possible imagined movements and auditory evoked potentials.

**Figure 2:**
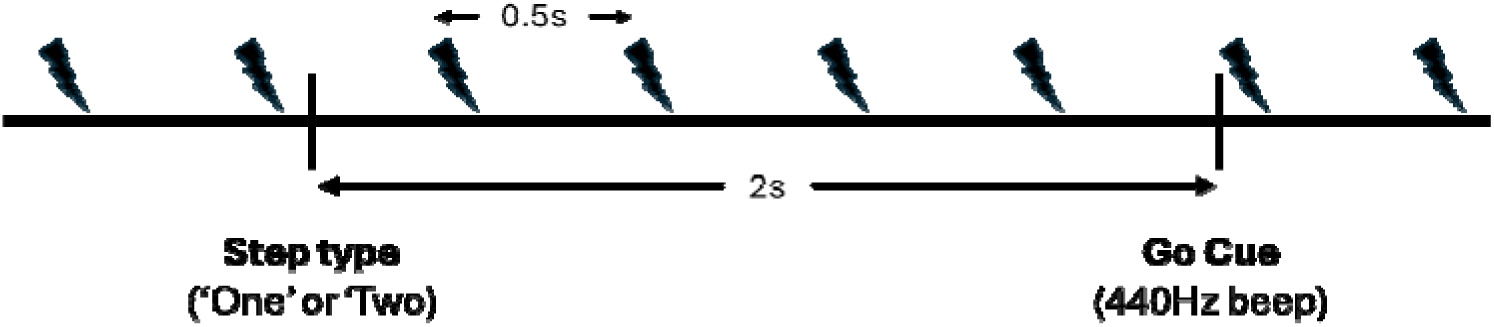
Electrical Stimulation and Auditory Cues. Lightning bolts represent the electrical stimulation. The timing of the cues remains the same for the CNV blocks.

### EEG Pre-Processing

The EEG data was consolidated into two files: one for the stepping trials and one for the standing trials. These files were imported into Python (version 2024.12.3) and analyzed using the mne library (version 1.7) (Larson et al., 2024). The following operations were done independently for these of the files in the order described below. Bad channels were removed, and their data was interpolated based on the remaining good channels. All electrodes were referenced to the average signal across all electrodes in their respective file. Notch filters at 60 Hz and its harmonics (60, 120, 180, 240, 300, and 360 Hz) were applied using a finite impulse response (FIR) filtering method to remove line noise. After notch filters’ application, Independent Component Analysis (ICA) with the fastICA algorithm was used for artifact removal to the greatest extent possible. Following artifact removal with ICAs, the data was filtered with a high-pass frequency of 0.1 Hz and a low-pass frequency of 40 Hz.

#### Epoching

To find the stimulation epochs, 0.05s prior to each electrical stimulation’s presentation until its onset was used as the epoch’s baseline period. The average value in each baseline period was subtracted from its respective epoch.

For creating each participant’s SEP epochs, the data from 0.05s prior to the electrical stimulation’s presentation (i.e., baseline) until 0.5 seconds after stimulation onset was extracted. Bad epochs were rejected to prevent them from contaminating the data in accordance with the following procedure.

First, epochs were also excluded if participants performed the task incorrectly or if there were deviations from the experimental protocol. This included instances where movement timing was inconsistent with task instructions (e.g., stepping on the warning cue), performing the wrong movement (e.g., stepping diagonally when instructed to step straight) or when other factors interfered (e.g., participant talking during the trials, phone ringing) with the expected execution of the task. By removing these epochs, it was ensured that only trials reflecting proper task execution and adherence to the study design were included in the final analysis, enhancing the reliability of the results. Removing these bad epochs resulted in an unequal number of good epochs across participants as the epoch’s exclusion criteria did not affect all participants similarly.

To increase signal to noise ratio in the good epochs remaining, 15% of the nosiest epochs were also removed. To identify and remove noisy epochs, an amplitude-based thresholding method was applied (Mumtaz et al., 2019). Epochs containing signals at any of the electrodes that exceeded a specific threshold were marked as noisy. This threshold was not fixed but was iteratively adjusted to ensure the removal of 15% of the total epochs, which were the noisiest segments of each participant’s data. By dynamically setting this threshold, only the most contaminated epochs were excluded, preserving as much clean data as possible in a participant-specific manner. If an electrode contributed to the removal of a significant portion of the epochs and its data deemed noisy with visual inspection, that electrode was removed, interpolated, and ICAs and epoch removal were then repeated with the removed electrode’s interpolated data.

### Evoked data

Clean epochs’ weighted averaging provided each condition’s evoked response. The SEPs were generated via the following two methods: In the pooled analysis, the epochs from the 4 electrical stimulations presented prior to each stepping condition (i.e., 4 stimulations prior to straight stepping and 4 stimulations prior to diagonal stepping) were combined to generate the SEPs. Additionally, SEPs were generated based on their timing relative to the auditory cues. To compute the evoked response while considering electrical stimulation’s order, the SEPs elicited from the 1^st^ electrical stimulation in the straight stepping condition, 1^st^ electrical stimulation in the straight diagonal stepping condition, SEPs elicited from the 2^nd^ electrical stimulation in the straight stepping condition, 2^nd^ electrical stimulation in the straight diagonal stepping condition, and so forth, were extracted and their weighted average was calculated. Pooled analysis aimed at reducing noise, while the ordered analysis optimized for analyzing the influences of different stages of motor preparation on sensory processing.

#### Source Localization

Good epochs and their evoked data then entered the following pipeline for source localization. The noise covariance was created based on the epoch’s base line period (from 0.05s to electrical stimulation’s presentation). A forward solution was created based on the Free Surfer average (fsaverage) brain’s source spaces, transformation file and boundary element method (Wu et al., 2018). Based on the forward solution, an inverse operator was created and the its inverse solution was applied to the evoked data using the eLORETA weighted minimum norm inverse solution (Pascual-Marqui, 2007). To manage the tradeoff between data fidelity and noise sensitivity in the inverse solution, the square of the regularization parameter λ (lambda) was used, i.e.:

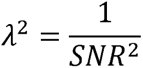

*Equation 1: λ^2^ was used as the noise sensitivity’s regularization parameter. SNR: Signal to Noise Ratio*

Based on the previous literature, a Signal to Noise Ratio (SNR) of 3 was deemed appropriate for the epoched data in such academic setting (Barzegaran et al., 2019; Beltrachini et al., 2021). The ‘*Automatic PARCellation, Destrieux Atlas, 2009, surface-based’*, was used to extract the mean regional activity for each sample frame across the evoked time window (Pascual-Marqui, 2007).

### Statistical Analysis

After the preprocessing steps above, participants’ evoked data and source localization data were exported to .csv files and imported to R Studio for statistical analysis. To evaluate condition related differences in the evoked data and the source localization data, a linear mixed-effects model was used. The model included Condition as a fixed effect and a random intercept for anonymized participants’ ID to account for within-subject variability due to repeated measures. This approach allowed for modeling individual baseline differences while properly accounting for the correlation between repeated measurements from the same participant. When multiple conditions were compared simultaneously (e.g., standing Vs. straight stepping, Vs. diagonal stepping), post hoc comparisons between conditions were conducted using estimated marginal means with Benjamini-Hochberg correction to control the false discovery rate. The linear mixed-effects models were tested for linearity, normal distribution, homoscedasticity, and points overly influencing the regression coefficients by examining their diagnostic plots. Multicollinearity was checked by the generalized variance inflation factor and observation’s independence by the Durbin-Watson test. No discrepancies to the model’s assumptions arose from examining these diagnostic plots and tests.

#### SEPs Analysis

SEPs’ statistical analysis focused on the Cz electrode in accordance with previous literature, and its proximity to cortical regions relevant for this study (S1 and M1). Previous literature also has established the first components for foot sole stimulation as P50 and N90 (Fabre et al., 2022; Mouchnino et al., 2015, 2017). Therefore, the P50 and N90 were respectively determined by extracting the maximum value occurring between 0.04s to 0.06s and the minimum value occurring between 0.08s – 0.10s at the Cz electrode. Then the difference between the P50 and N90 components provided a peak to peak amplitude. These SEP measures (P50, N90, and peak to peak amplitudes) were compared by pooling the SEP measures in each stepping condition, and by analyzing the evoked response based on the order in which the electrical stimulation was presented at. The pooled analysis optimizes for increasing signal to noise ratio, reflecting a general task-level effect on SEP amplitude across repeated stimulations. The ordered analysis aimed at capturing the potential influences of the preparatory processes’ stage on sensory processing.

In the pooled analysis, the evoked SEP measures from all four stimulations between the Warning and Go cues, along with the measures from all the stimulations in the Standing condition were pooled into a single dataset. Each participant’s evoked SEP measurement formed an observation. The pooled SEP measures were compared across the three task conditions (Standing, straight stepping, and diagonal Stepping), and post hoc comparisons between conditions were conducted using estimated marginal means with Benjamini-Hochberg correction to control the false discovery rate. For the ordered analysis, the SEP measures were sorted based on their timing relative to the auditory cues (i.e., 1^st^, 2^nd^, 3^rd^, 4^th^ SEPs in the planning of each Stepping condition). Again, each participant’s evoked SEP measure formed an observation. The ordered SEP measures were only compared between the Stepping tasks (i.e., straight Vs. diagonal) because the preparatory stage’s influences do not affect the Standing condition where movement planning was absent.

#### SEP Source Localization Analysis

The source activity in each extracted region was averaged according to the time windows used for finding the SEP components’ peak amplitudes (i.e., region’s mean during 0.04s – 0.06s for P50, and 0.08s – 0.1s for the N90). The following process, following literature’s guidelines, was used to parse out relevant cortical regions with sufficient cortical activation (Barzegaran et al., 2019; Beltrachini et al., 2021). First, a threshold was calculated by determining 80% of the activation amplitude in the region maximally activated for each component for each participant. Regions that met the 80% threshold in at least 50% of the participants were kept for further analysis. The mean activity for the different SEP components in the kept regions were compared between the stepping conditions using the mixed effects linear model. Similar to the SEPs, a pooled analysis was done to explore source contributors to the SEPs in a general manner with high signal to noise ratio level, while an ordered analysis captured preparatory stage influences.

## Results

### Pooled Analysis

In the pooled analysis, the mixed effect linear model showed a significant difference in the peak-to-peak amplitude between Standing condition and straight stepping (P = 0.041), Standing Condition and diagonal stepping (P = 0.026), but no significant difference between the straight and diagonal Stepping conditions (P = 0.586).

Figure 3 shows the SEP traces at the Cz electrode for the pooled analysis for the stepping and standing conditions.

**Figure 3:**
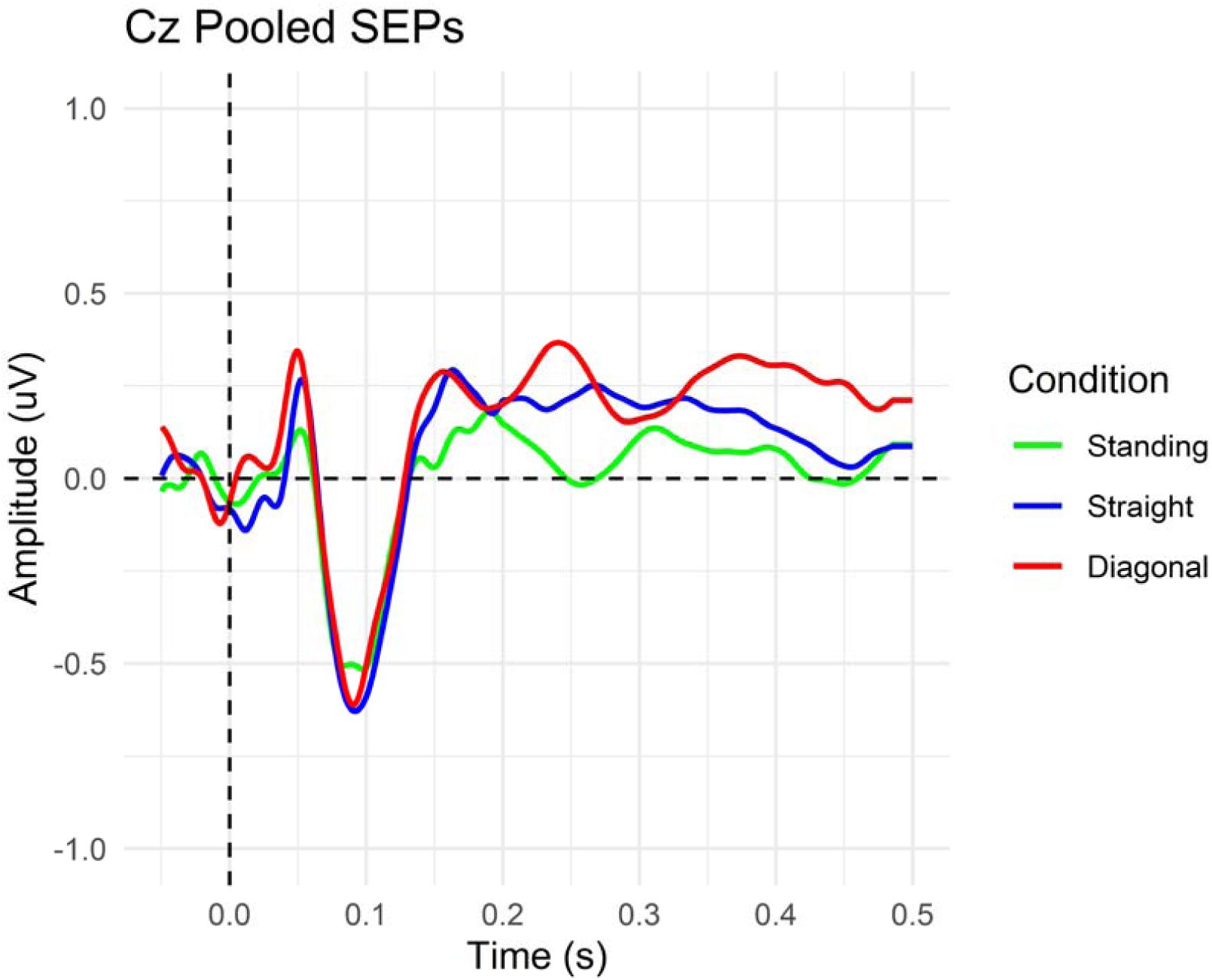
Pooled SEP Evoked Traces at the Cz Electrode

Table 1 summarizes the P values between the conditions for the first SEP components (P50 and N90) and their peak to peak measure in the pooled analysis. A significant difference for the peak to peak measure between standing and either of the stepping conditions exists.

**Table 1:** Pooled SEP Components’ statistical values from the mixed effects model.

| **Statistical Values for SEP Components in the Pooled Analysis** |  |  |  |  |
| --- | --- | --- | --- | --- |
| **P50** |  |  |  |  |
| **Compared Conditions** | **Estimate** | **Standard Error** | **t Value** | **P Value** |
| Standing vs. Straight | -0.099 | 0.103 | 0.960 | 0.364 |
| Standing vs. Diagonal | -0.193 | 0.103 | 1.875 | 0.198 |
| Straight vs. Diagonal | -0.094 | 0.103 | 0.914 | 0.364 |
| **N90** |  |  |  |  |
| **Compared Conditions** | **Estimate** | **Standard Error** | **t Value** | **P Value** |
| Standing vs. Straight | 0.158 | 0.118 | 1.344 | 0.419 |
| Standing vs. Diagonal | 0.128 | 0.118 | 1.092 | 0.419 |
| Straight vs. Diagonal | -0.030 | 0.118 | 0.252 | 0.802 |
| **Peak to Peak** |  |  |  |  |
| **Compared Conditions** | **Estimate** | **Standard Error** | **t Value** | **P Value** |
| Standing vs. Straight | -0.257 | 0.118 | 2.173 | 0.041 |
| Standing vs. Diagonal | -0.322 | 0.118 | 2.720 | 0.026 |
| Straight vs. Diagonal | -0.065 | 0.118 | 0.547 | 0.586 |

Figure 4 presents the distribution of the peak to peak measurements across the conditions in the pooled analysis.

**Figure 4:**
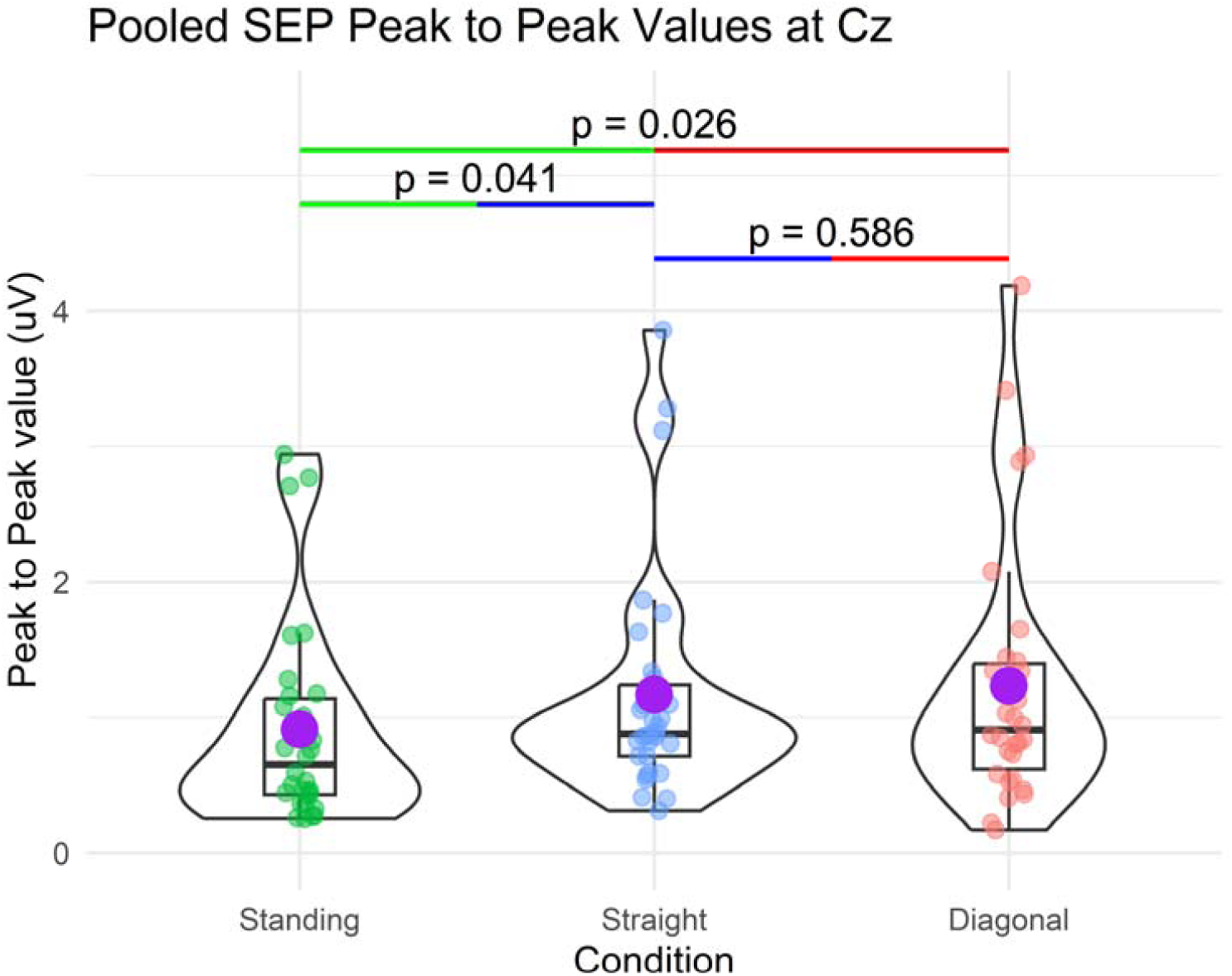
Distribution of the peak-to-peak values from Pooled SEPs and the Standing SEPs. The line in the box plot represents quartile ranges with line inside the box representing the median. The purple dot represents the group’s mean.

### Ordered Analysis

Analyzing the SEPs based on the order in which they were presented highlighted a significant difference between the straight stepping and diagonal stepping conditions for the 1st electrical stimulation (P = 0.009), which was presented at 0.5s after the presentation of the Warning cue. There were no significant differences for the remainder of the SEPs. The significant difference in the peak-to-peak amplitudes for the 1^st^ SEP suggests this information is amplified at the early stages of motor preparation. Additionally, this result suggests the different stages of motor preparation may influence the incoming somatosensory information differently.

Figure 5 shows the SEP traces at the Cz electrode for the SEP from the 1^st^ electrical stimulation in the stepping conditions.

**Figure 5:**
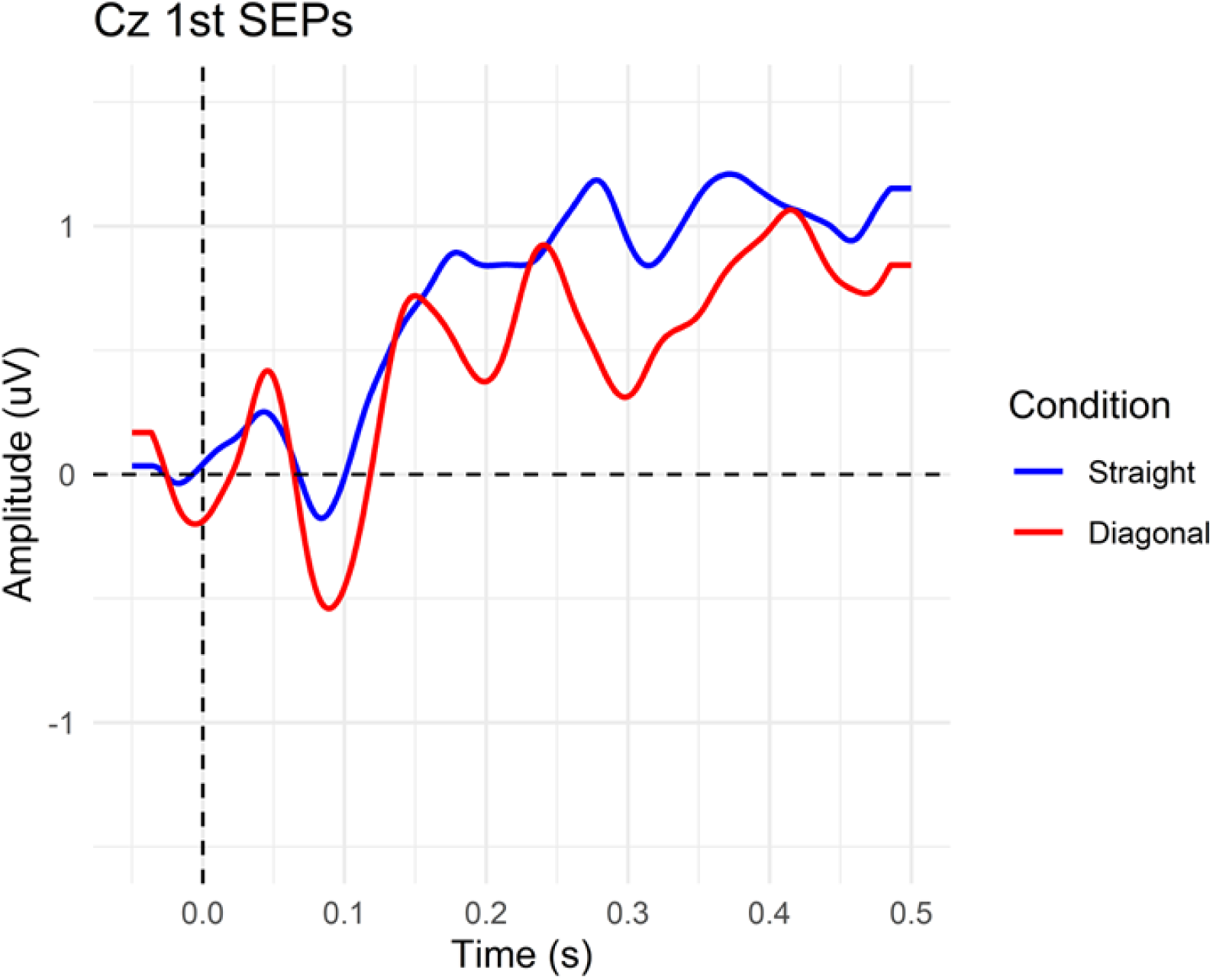
1st SEP Evoked Trace at the Cz Electrode

Table 2 summarizes the P values between the stepping conditions for the first SEP components (P50 and N90) and their peak to peak measure for the ordered SEPs at the Cz electrode. A significant difference for the peak to peak measure between the stepping types is present for the 1^st^ SEP.

**Table 2:**
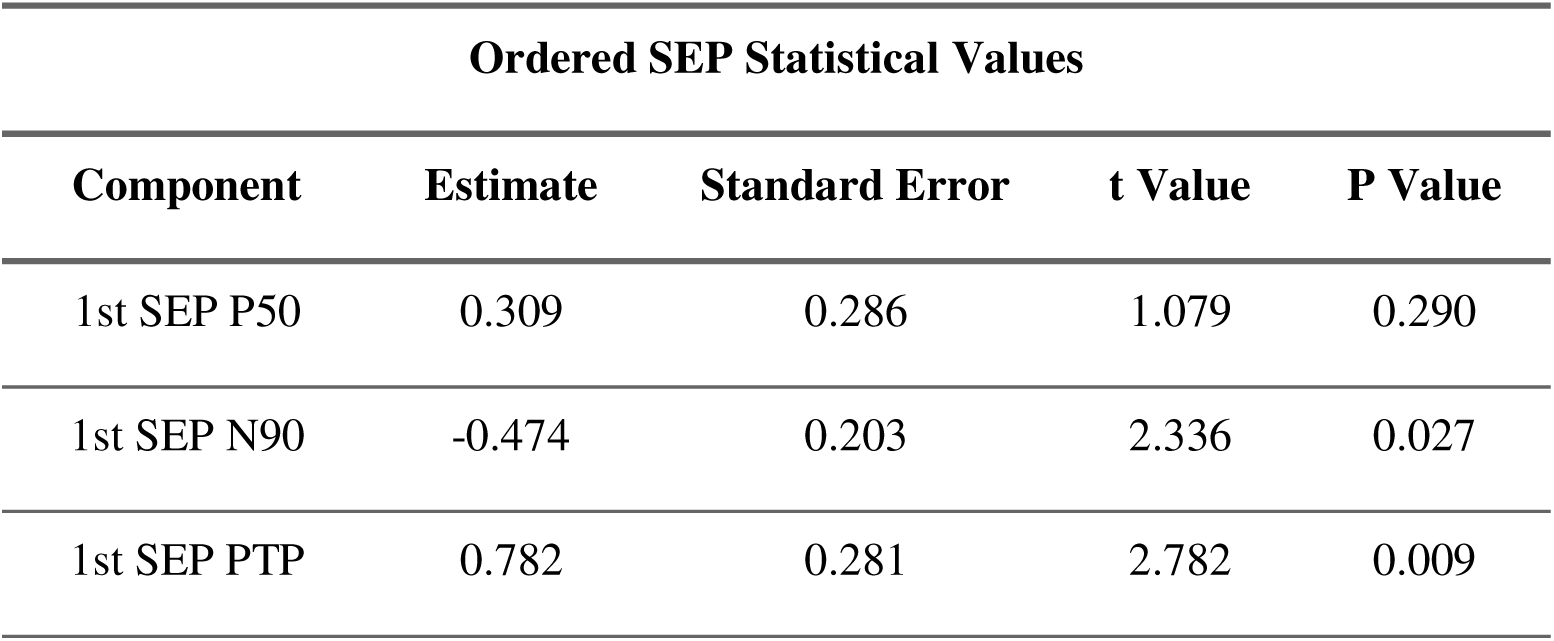
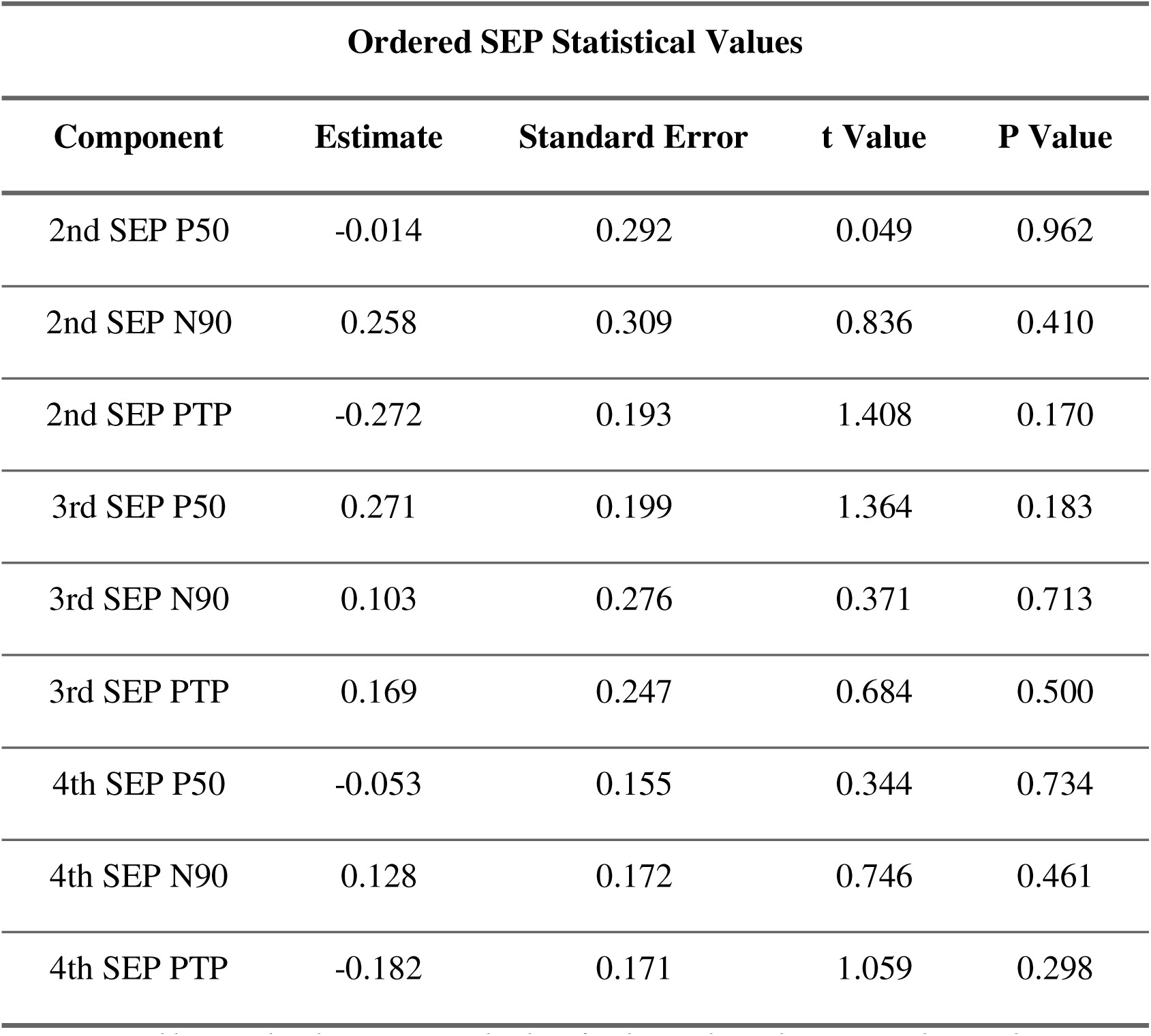
Ordered SEP statistical values for the Cz electrode. PTP: Peak to Peak.

| Ordered SEP Statistical Values |  |  |  |  |
| --- | --- | --- | --- | --- |
| Component | Estimate | Standard Error | t Value | P Value |
| 1st SEP P50 | 0.309 | 0.286 | 1.079 | 0.290 |
| 1st SEP N90 | -0.474 | 0.203 | 2.336 | 0.027 |
| 1st SEP PTP | 0.782 | 0.281 | 2.782 | 0.009 |
| Component | Estimate | Standard Error | t Value | P Value |
| 2nd SEP P50 | -0.014 | 0.292 | 0.049 | 0.962 |
| 2nd SEP N90 | 0.258 | 0.309 | 0.836 | 0.410 |
| 2nd SEP PTP | -0.272 | 0.193 | 1.408 | 0.170 |
| 3rd SEP P50 | 0.271 | 0.199 | 1.364 | 0.183 |
| 3rd SEP N90 | 0.103 | 0.276 | 0.371 | 0.713 |
| 3rd SEP PTP | 0.169 | 0.247 | 0.684 | 0.500 |
| 4th SEP P50 | -0.053 | 0.155 | 0.344 | 0.734 |
| 4th SEP N90 | 0.128 | 0.172 | 0.746 | 0.461 |
| 4th SEP PTP | -0.182 | 0.171 | 1.059 | 0.298 |

Figure 6 presents the distribution of the peak to peak measurements between the stepping conditions for the 1^st^ SEP.

**Figure 6:**
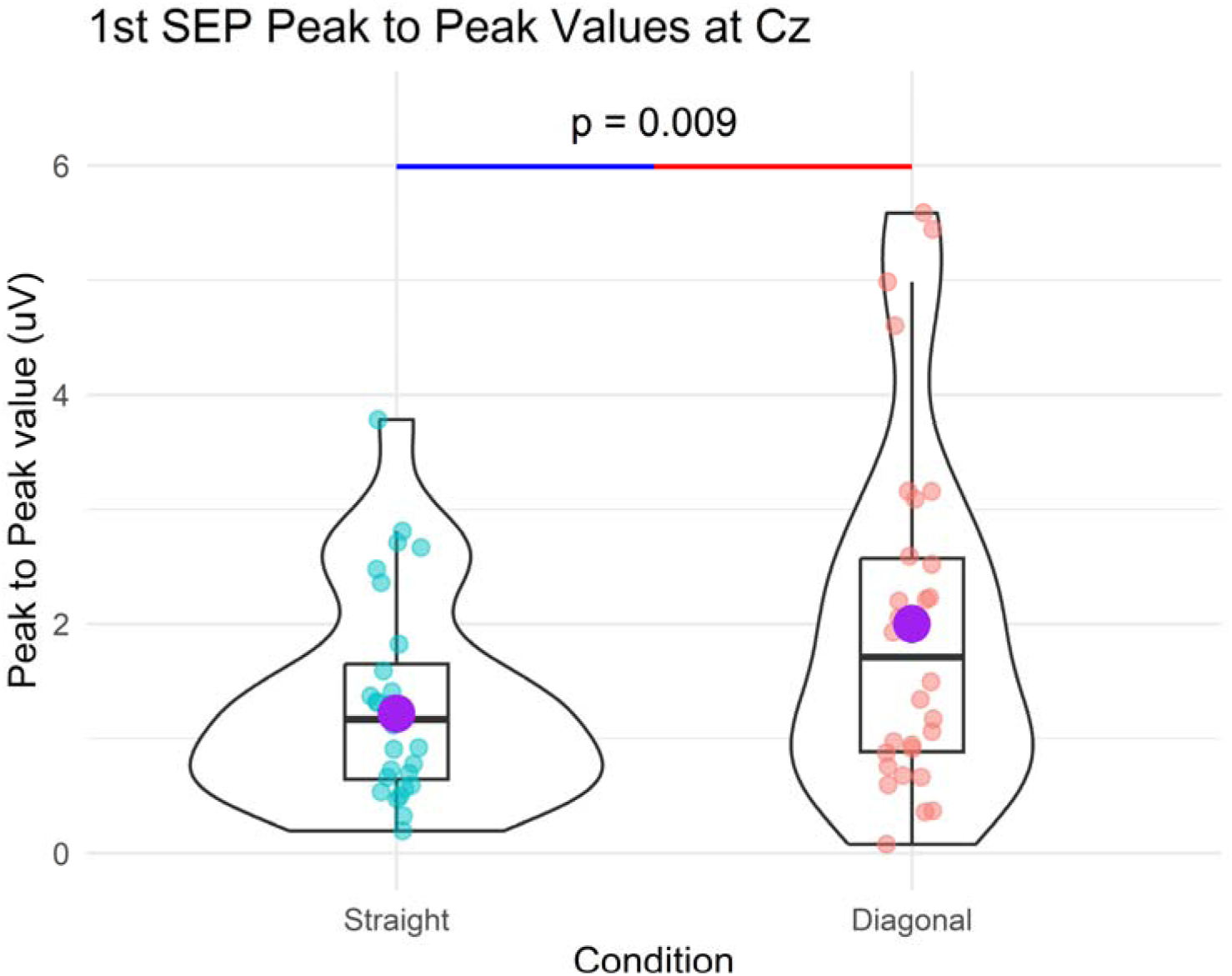
1st SEP peak-to-peak violin plot for the Cz electrode. The line in the box plot represents quartile ranges with line inside the box representing the median. The purple dot represents the group’s mean.

### Source Localization

#### Pooled Source Localization

Source Localization Analysis for the Pooled SEPs highlighted the following cortical regions as being significantly more active in both Stepping conditions compared to the Standing Conditions at the N90’s time window: Paracentral Gyrus & Sulcus, Inferior Parietal Angular Gyrus, and Superior Parietal Gyrus. No significant differences were found for the P50 component, or between the Stepping Conditions. None of the cortical regions showed a decreased activity in the Stepping condition compared to the Standing condition at any of the SEP components.

Table 4 summarizes statistical measurements for the cortical regions significantly more active between conditions in the pooled analysis, at the N90 time frame. The conditions are compared in the order listed in the table, indicating the direction of the difference in the t-values. For example, a positive t-value for Standing vs. straight highlights increased cortical activation in the straight condition compared to Standing, for the region listed in the table. Note these cortical regions are significantly more activated in both stepping conditions compared to standing, but they are not significantly different between the stepping conditions.

**Table 3:**
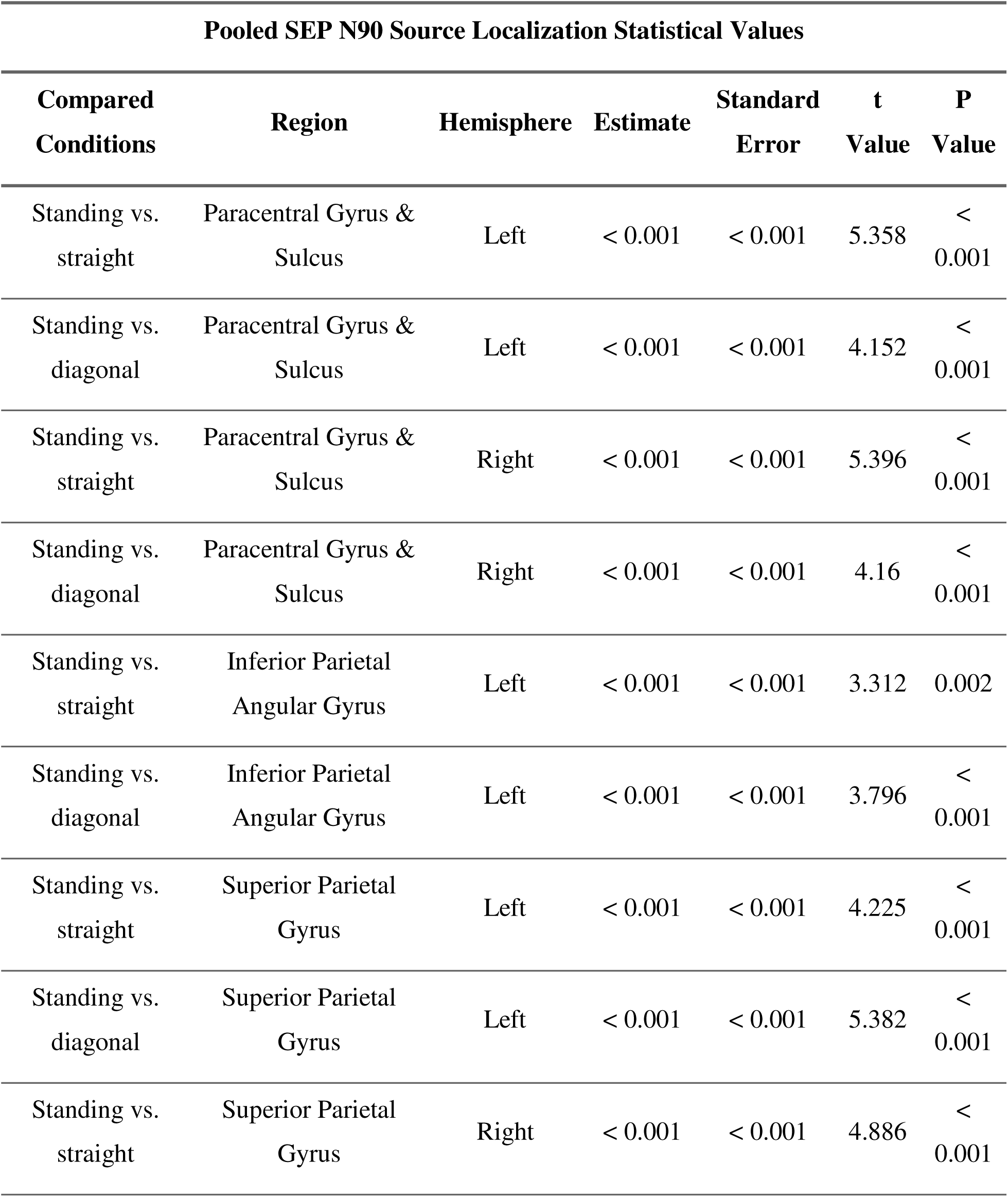
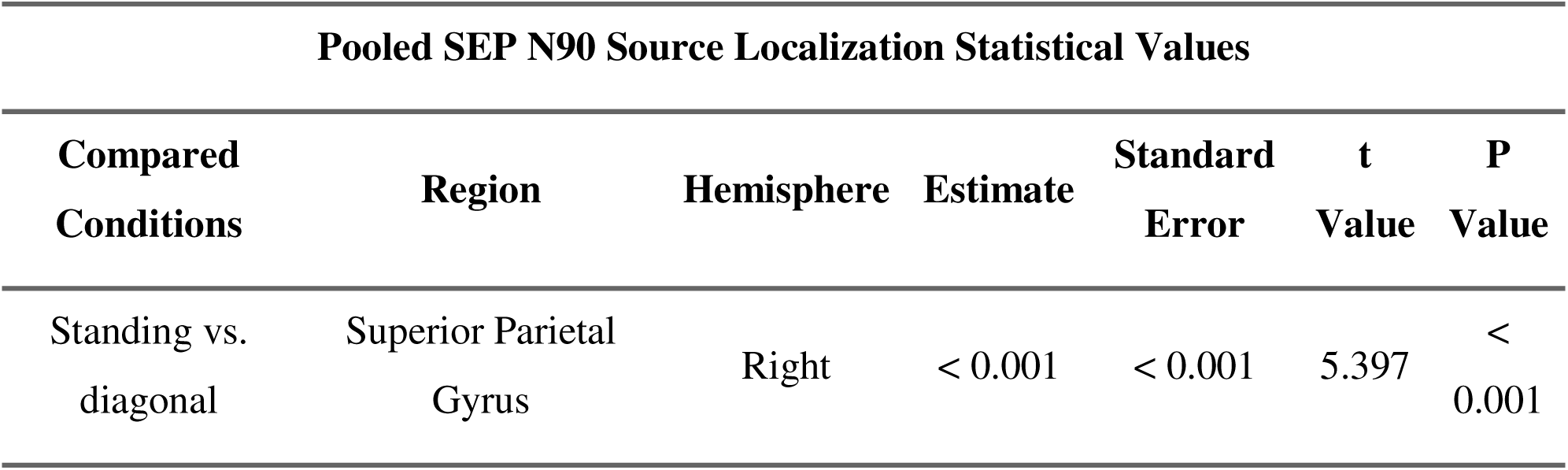
Significantly more active regions statistical data for the N90 time frame from SEP’s pooled source localization analysis.

Figure 8 locates the cortical regions significantly more active in the stepping conditions compared to standing.

**Figure 7:**
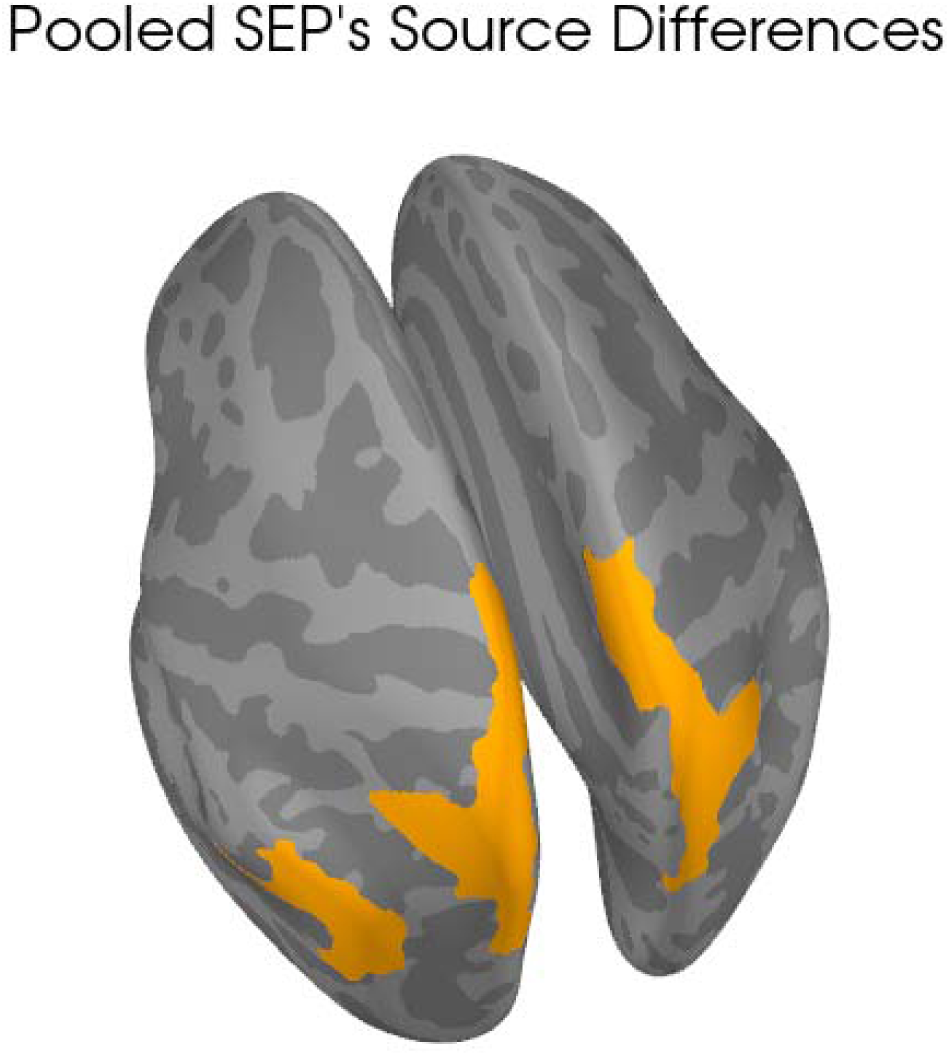
Significantly different cortical regions illustrated in yellow. These areas were more active for the N90 time frame between the Stepping conditions compared to Standing, but not between Stepping Conditions.

#### Ordered Source Localization

Ordered Source Localization highlighted the following region as being significantly more active in the diagonal stepping condition at the N90’s time window at the 1st, and 4th SEPs: Paracentral Gyrus & Sulcus. No significant differences were found for other orders of the SEPs (2nd and 3rd) or other components (P50).

Table 4 summarizes statistical measurements for the cortical regions significantly more active between conditions in the ordered analysis, at the N90 time frame. straight stepping is compared to diagonal stepping. Hence, a positive t-value indicates increased cortical activation in diagonal stepping compared to straight stepping.

**Table 4:**
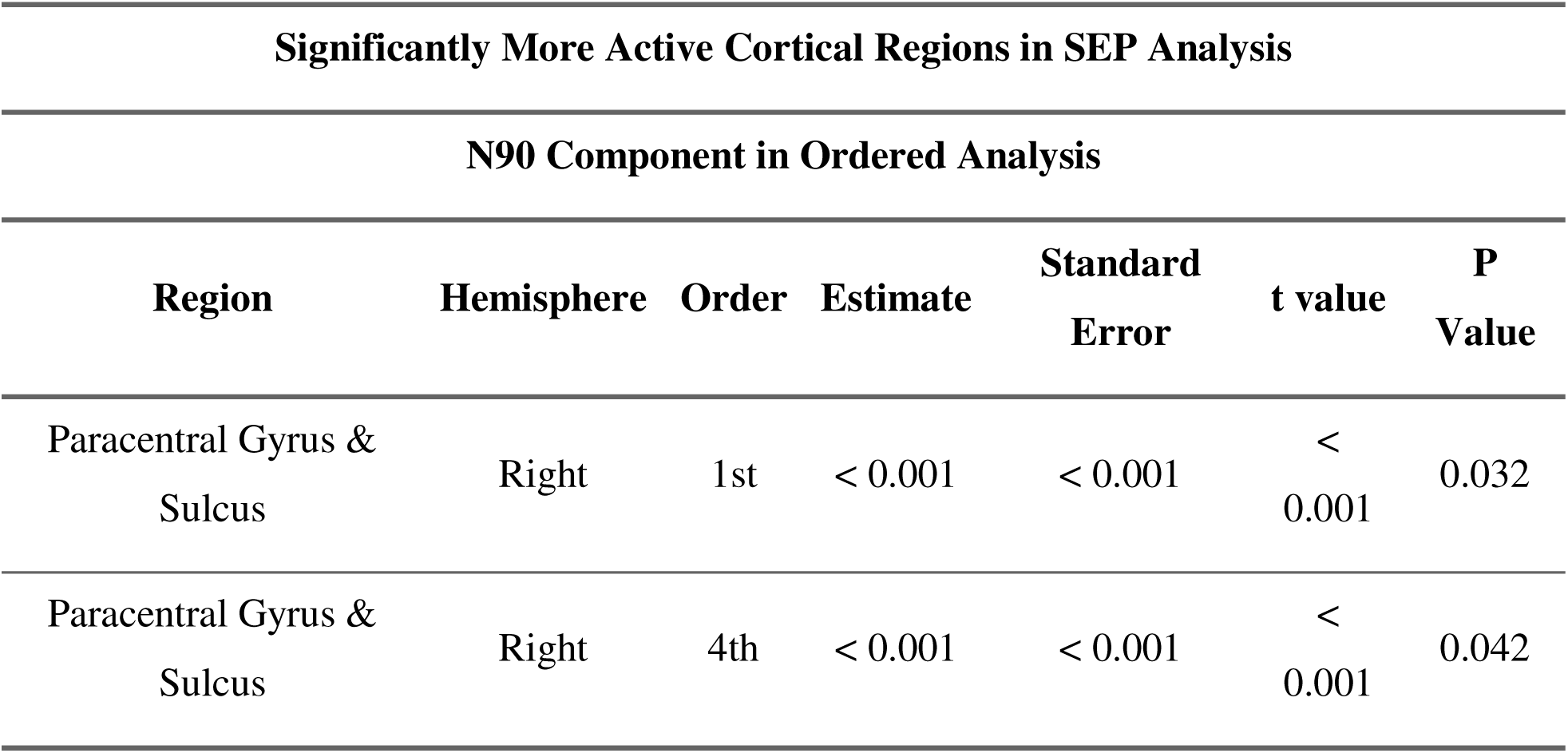
Significantly more active regions statistical data for the N90 time frame from ordered eLORETA analysis between straight and diagonal stepping.

Figure 8 shows right Paracentral Gyrus & Sulcus, the cortical region significantly more activated in the diagonal stepping condition, compared to straight stepping, during the N90 time frame in the 1^st^ and 4^th^ electrical stimuli. Paracentral Gyrus & Sulcus is near the S1’s leg region, and the right hemisphere is contralateral to the electrical stimulation site.

**Figure 8:**
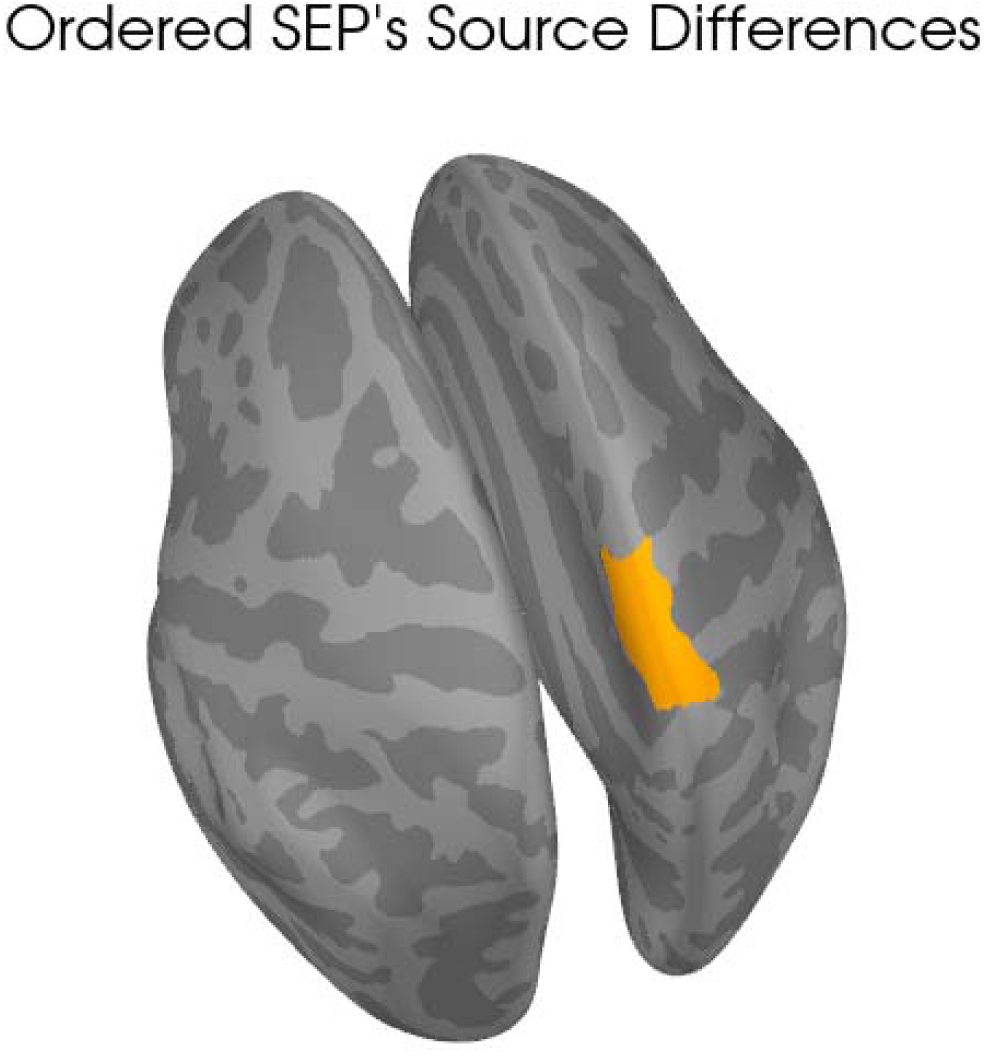
Significantly different Cortical Regions illustrated in yellow. These areas were more active for the N90 time frame between the Stepping conditions in the Ordered Analysis for the 1st and 4th SEPs

## Discussion

The different levels of postural complexity may be influencing the foot sole cutaneous information. Given the SEPs highlighting brain’s response to the incoming foot sole cutaneous information, responding to the task’s balance demands via postural modulations may be affecting the measured SEPs. Anticipatory selective attention mechanisms, via pre-frontal cortex’s control over the thalamus gating may be a cortical mechanism influencing SEPs amplifications in tasks with a higher postural complexity (Ariani et al., 2022). M1’s impact over the S1 and the thalamus may be another cortical mechanism influencing the measured SEPs (Gómez et al., 2021).

### Postural Complexity

In the tasks performed in this study, the postural complexity differs in response to task’s balance requirements. Balance and postural requirements in tasks performed in this study progress in their complexity in the following order: standing, straight stepping, and diagonal stepping.

Maintaining balance in standing is simplest and requires the least postural modulations from the tasks mentioned above. There is minimal threat to balance in standing (Fabre et al., 2021; O’Connor & Kuo, 2009). CoM is situated well within BoS, requiring simple muscle activation patterns for postural stability (Fabre et al., 2021; O’Connor & Kuo, 2009). Straight stepping and diagonal stepping are both more complex than standing. In stepping, the body weight must be propelled forward, and the swinging limb accurately placed to re-establish a stable BoS (Bancroft & Day, 2016). Creating the momentum required for thrusting the body forward, controlling this forward momentum, and coordinating the leg swing to land at an optimal location for regaining balance requires intricate control (Bancroft & Day, 2016; Lyon & Day, 1997). APAs, generated before the swing foot leaves the ground, shift load onto the stance limb, produce the forward body thrust, and preset the swing leg’s trajectory and timing (Alfuth & Rosenbaum, 2012). Their precise scaling relies on foot-sole cutaneous input, which signals pressure distribution and load location needed to plan these adjustments feed-forward (Alfuth & Rosenbaum, 2012). Enhancing this somatosensory channel would therefore facilitate the precise timing and scaling of APAs needed to initiate and complete the step safely and efficiently.

In the frontal plane, the key variable at swing-foot lift-off is how far the CoM is offset from the stance foot’s BoS as this offset sets the lateral momentum for the step (Lyon & Day, 1997). For straight stepping, APAs shift the CoM laterally toward (often just to the edge of) the stance foot’s BoS to create the small but necessary lateral impulse while keeping the CoM largely supported (Assaiante et al., 2000). For diagonal stepping, the APAs push this shift further: the CoM is moved beyond the stance BoS, creating a larger lateral offset (Assaiante et al., 2000).

Once the swing foot leaves the ground, this offset lets the body “fall” sideways around the stance ankle, generating the required lateral component of momentum (Bancroft & Day, 2016; Chen & King, 2024; Lyon & Day, 1997). Because the CoM starts farther from the stance BoS in diagonal steps, the sideways acceleration is greater, making timely foot placement more critical and increasing postural control demands relative to straight stepping (Bancroft & Day, 2016).

Diagonal stepping forces APAs to generate and control a larger lateral momentum, so they must be planned and scaled more precisely than in straight stepping. The increased intricacy in planning the APAs in diagonal stepping makes foot-sole cutaneous input more critical for diagonal stepping than in straight stepping.

Electrical stimulation of the foot sole provides a standardized proxy for plantar cutaneous input; the resulting SEPs are therefore a controlled readout of the brain’s response to that specific somatosensory channel (Kappenman & Luck, 2011). Because APAs in more balance-demanding tasks depend more heavily on accurate plantar feedback, the information foot sole cutaneous receptors are amplified in tasks with a higher balance demand compared to simpler balance demands. Therefore, larger SEPs are observed in the stepping conditions compared to standing, and larger SEPs in diagonal stepping compared to straight stepping.

### Cortical Mechanisms

Several cortical mechanisms may be involved in modulating the SEPs elicited from the foot sole stimulation. The thalamic gating may be controlled via the prefrontal cortex, the M1 may be influencing thalamic and S1 activity (Alonan & Brown, 2002; Carmona et al., 2023; Gómez et al., 2021). In the earlier phases of preparing to take a step, anticipatory selective attention mechanism may be involved in directing attention towards relevant foot sole information (ElShafei et al., 2018). Later in the preparatory phase, this somatosensory information may be integrated into the motor plan (Gaillard, 1977).

#### Prefrontal cortex and Thalamic gating

The prefrontal cortex can bias thalamic relay nuclei in a top-down manner, opening the gate for task-relevant input and damping irrelevant signals (Aydin et al., 2022). Under higher balance demands, when precise plantar feedback is needed to scale APAs, this control could amplify foot sole afferents reaching S1, producing larger SEPs. When demands are lower, the gate can remain tighter, limiting plantar input and attenuating SEP amplitude.

If prefrontal cortex driven thalamic gating amplifies plantar afferents when balance demands rise, it would yield a graded SEP pattern across our tasks (Allison, 1962; Callier et al., 2019). During standing, limited postural challenge keeps the “gate” relatively constrained, so fewer foot-sole signals reach S1 and SEP amplitudes remain small. Initiating a step requires more precise APAs and thus opening of the gate, increasing SEP size. Diagonal stepping imposes greater postural complexity than straight stepping, prompting further thalamic gate opening and increased amplification of this somatosensory information. Increased amplification somatosensory amplification leads to the largest proportion of plantar input reaching the cortex, producing SEPs such that Diagonal > Straight > Standing.

#### M1’s influence

M1 may also be involved in modulating the incoming sensory information, by influencing thalamic gating and S1 activity (Canedo, 1997; Gómez et al., 2021). By sending a copy of the expected sensory outcomes of a movement, M1 can reduce the activity in the S1 caused by these expected sensory outcomes, allowing unexpected sensory outcomes, such as errors or deviations from the prepared movement plan, to be identified and responded efficiently (Gómez et al., 2021; Wurtz & Sommer, 2004). For tasks where the sensory outcome of the movement cannot be predicted with a high level of certainty, M1 does not inhibit S1 (Altermatt et al., 2023; Canedo, 1997; Wurtz & Sommer, 2004). Since the weight shifts in the APAs must be performed intricately to deal with the task’s balance demands, M1 may not be inhibiting the expected sensory outcomes from the APAs. This inhibition may be further reduced for the tasks with higher balance demands (i.e., diagonal stepping compared to straight stepping), to provide additional information for APA’s planning (Goodworth & Peterka, 2012).

#### Preparation’s different stages

Different cortical mechanisms may be involved in the movement preparation’s different stages (Fattapposta et al., 2024; Gaillard, 1977). Immediately after the warning cue in our two-cue paradigm, anticipatory selective attention is deployed: prefrontal networks bias thalamo-cortical processing toward sensory channels that will matter for the forthcoming action (Aydin et al., 2022; Goldman-Rakic, 2007). This early, top-down focus raises the gain on plantar afferents, so electrical foot-sole stimulation delivered near the 1^st^ cue evokes an enlarged SEP. Because diagonal stepping depends more critically on precise lateral load feedback than straight stepping, the attentional up-weighting of foot-sole input is stronger, producing a larger first-interval SEP in Diagonal > Straight conditions. As the go cue nears, the brain shifts from monitoring sensations to planning the movement; parietal cortex and S1 then blend foot-sole input into the motor plan (Brunia, 1993; Kawato, 1999; Sober & Sabes, 2005). This late-stage sensorimotor integration boosts SEP amplitude for stepping versus standing, and the added lateral-balance demands of diagonal stepping require still finer integration (Brunia, 1993; Thoenissen et al., 2002). Consequently, the foot-sole stimulus delivered closest to movement onset (4th pulse) evokes the strongest contralateral S1 response in the Diagonal > Straight comparison.

## Conclusion

This study demonstrates that motor tasks with varying postural demands elicit distinct patterns of cortical activation, reflecting their unique balance constraints. The presence of APAs enhances cutaneous sensory information originating from the stance foot sole. More intricate APAs in diagonal stepping, compared to straight stepping, lead to further amplification of stance foot sole’s cutaneous information; however, only during periods of anticipatory attention shift to the upcoming motor task and when motor preparation is occurring.

